# muDM: A Queryable Multi-Resolution Data Model for Vector Geometry in Spatial Bioimaging

**DOI:** 10.64898/2026.10.16.759449

**Authors:** Bengt Ljungquist, Nathan Hotaling, Nick Schaub

**Affiliations:** NovaGen Research Fund (NGRF), Rockville, MD, USA; National Center for Advancing Translational Sciences (NCATS), National Institutes of Health (NIH), Bethesda, MD, USA

**Keywords:** spatial bioimaging, vector data model, data linking, level of detail, multi-resolution pyramid, connectomics, spatial transcriptomics, machine learning, FAIR data

## Abstract

Advancements in spatial bioimaging analytics locate regions of interest as mask or vector geometries from raster images that are associated with molecular measurements, connectivity, and anatomical labels. A major challenge is to connect and jointly query these regions and their associated metadata at scale. The MicroData Model (muDM) is a spatial data model for associating 2D and 3D vector geometry with its annotated records, including identities, measurements, and hierarchies, as a precomputed multi-resolution pyramid. The data model has built-in integrations that emit target formats depending on use case, for consumption by machine learning and visualization applications. An open-source Python implementation is available on the Python Package Index, with a public catalog of over 100 datasets. In two *Drosophila* connectomes, the coarsest level retained at least 87% of finest-level neuron-typing accuracy with about 1% of the stored geometry. Linking synapse counts to the meshes improved accuracy beyond geometry alone.

## Introduction

Spatial bioimaging captures vector-based geometries from information-rich raster images, including segmentation-derived cell outlines and anatomical regions as well as surface meshes. Advances in image analysis allow processing at scale, for example tens of thousands of connectome neurons ^1^, and spatial transcriptomics where there are point-cloud representations of millions of transcript detections ^2^. These geometric objects connect different types of measurements and spatial hierarchies. A cell outline associates an image area with an expression profile, while a neuronal mesh links anatomical form to connectivity; nuclei are contained within cells, and cells within tissue regions. Analyses need to not only associate measurements with regions but also traverse hierarchical relationships, including, for example, retrieving image-derived features, transcripts, or morphological properties for structures located in a specified region. A dense transcript field may contain hundreds of megabytes of data. Although raster imagery of this scale can use chunked multi-resolution formats and expression matrices can use columnar storage, the vector geometry linking those data lacks an equivalent that supports analytic queries. No standard format or data model specifies how this geometry is stored across multiple resolutions, together with the records it annotates, and enables ordinary queries to retrieve the level required by a task. As a result, projects repeatedly re-tile or re-encode geometry, linking data takes significant time and effort, analyses often use full-detail data when a lower-resolution representation would suffice, and 2D and 3D annotations remain in separate, incompatible software stacks.

Current formats address part of this problem, but none provides the full combination. SpatialData creates pyramids for images and label masks, while retaining points and shapes at one resolution ^3^; the OME-NGFF specification behind OME-Zarr supports multiscale rasters but defines no vector-geometry type ^4^; and Celldega places single-resolution Parquet transcript and cell-boundary tiles behind an image pyramid^5^. This remains true even in resources explicitly intended for machine learning (ML). The spatial-proteomics dataset resource spora, packaged for foundation-model training, represents cell segmentation as single-resolution raster masks ^6,7^, and the spatialproteomics toolbox similarly stores cells as labeled raster images^8^. Existing vector and mesh level-of-detail pyramids target visualization more than analytical use: Open Geospatial Consortium (OGC) 3D Tiles ^9^ and Potree ^10^ deliver geometry to GPUs, while Mapbox Vector Tiles ^11^ encode 2D map features for the web. However, all require specialized decoders and yield scene graphs or rendering buffers instead of filterable columnar arrays. TileDB-SOMA combines vector geometry, columnar storage, and a resolution pyramid, but applies the pyramid only to raster imagery and leaves point and polygon tables at a single resolution ^12^. GeoParquet ^13^ makes geometry queryable but, like the related GeoArrow specification ^14^, defines neither levels of detail nor an indexed surface-mesh type. The absent capability is a queryable pyramid for vector geometry that unifies 2D and 3D, while retaining the biological meaning of its stored features.

The MicroData Model (muDM) addresses this need. It combines a spatial data model, in which every feature includes geometry, measurements, a link to its parent structure, and references to the external record it annotates, with a tiling workflow that computes a multi-resolution geometry pyramid in advance. At every pyramid level, data are emitted both as a columnar representation that can be queried directly and in standard rendering formats. A single ingest therefore supports visualization and analysis, and resolution is selected through a query instead of another encoding step (Fig. 1).

**Fig. 1.**
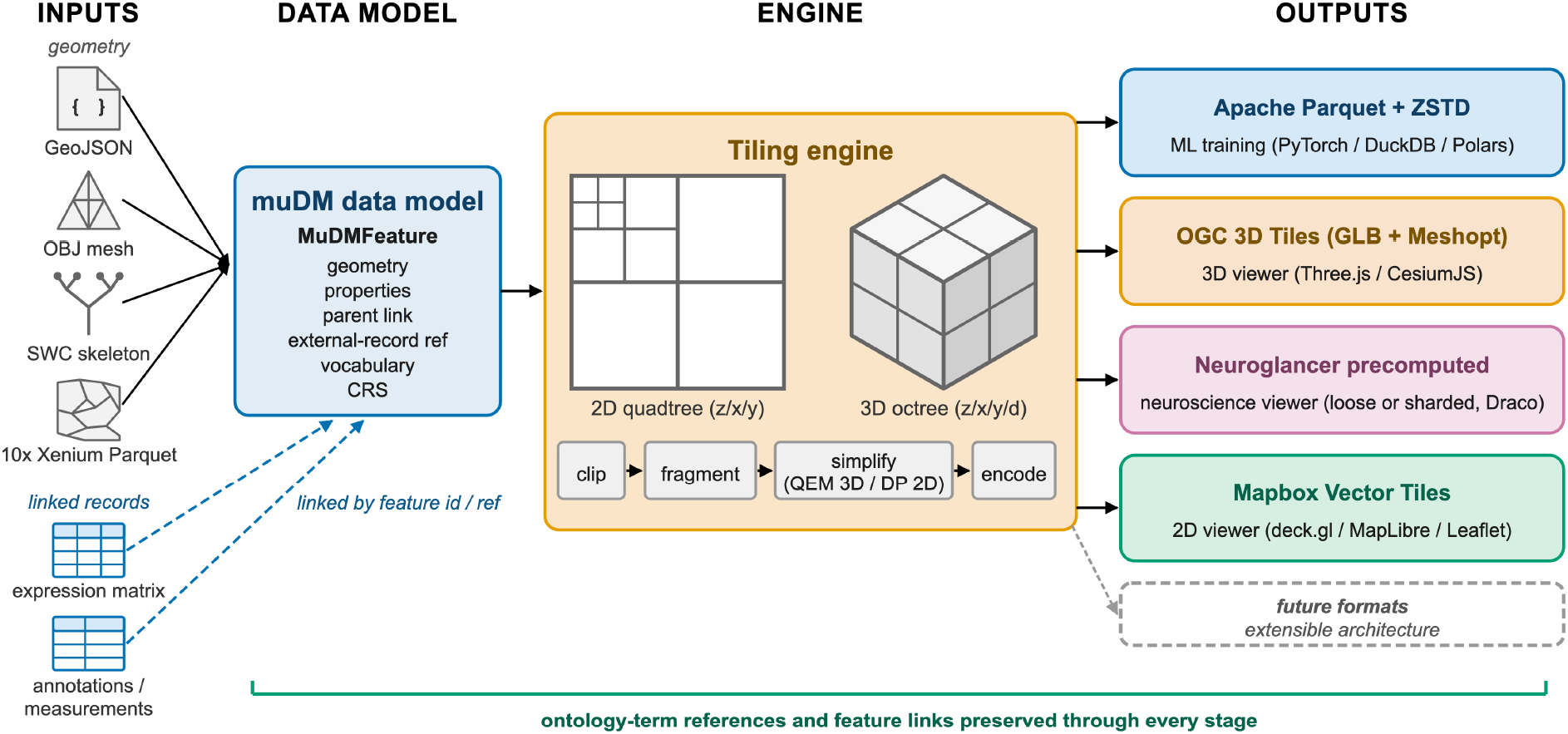
muDM multi-format tiling pipeline. Geometry from GeoJSON, meshes, skeletons, and vendor tables, together with associated records such as expression matrices, annotations, and measurements, is ingested into a unified data model. Features include geometry, properties, parent relationship, external-record reference, and controlled-vocabulary terms. The tiling engine spatially indexes geometry using a 2D quadtree or 3D octree, then generates a precomputed multi-resolution pyramid. Every level is output both as a queryable columnar store, Apache Parquet with Zstandard (ZSTD) compression, and in standard visualization formats: OGC 3D Tiles, Neuroglancer precomputed meshes, and Mapbox Vector Tiles. Feature relationships and ontology-term references remain intact throughout the pipeline.

We additionally provide a public catalog, organized using the muDM data model, and containing more than 100 datasets across connectomics, multiplexed tissue imaging, and spatial transcriptomics. It is available through browser-based reference implementation viewers and is also structured for programmatic and agent-driven access ^15^. Experiments on *Drosophila* connectomes quantify what the linked, multi-resolution store enables. A classifier given synapse counts linked to each neuron mesh outperformed one given geometry alone, and lower-resolution pyramid levels preserved most neuron-typing accuracy while reducing stored geometry by up to two orders of magnitude. The 3D mesh-serving approach extends existing standards. muDM is available as open source under an MIT license, including the mudm data model and the mudm-tools pipeline with web viewers for both 2D and 3D.

## Results

### muDM links 2D and 3D geometry to its records and hierarchy at every resolution

Every muDM feature carries its geometry together with the records that describe it, as data are interlinked. The model is derived from the GeoJSON feature model ^16^, adding microscopy-specific 3D geometry types, physical coordinate systems, and controlled-vocabulary metadata that provides the capability to associate every feature with an ontology term. Beyond these, each feature holds an unrestricted measurement property set, as well as a pointer to a parent structure (for example, a nucleus to its cell or a cell to its tissue region), and a pointer to an external record or repository, such as the expression matrix containing a cell’s readout. A 2D polygon outlining a cell boundary and a 3D neuron mesh therefore both use the same model, identifiers, semantic labels, structure, and links.

The pyramid is column-oriented, so only the required columns and row groups at the specified resolution are read. We may, for example, query a specific level of detail and filter on spatial region, and limit to a range of objects, all read from storage directly without scene-graph decoding (Supplementary Note 4). Filters can also target metadata fields, feature identifiers, and tag values. Geometries are stored as packed arrays that PyArrow exposes without copying as NumPy or PyTorch ^17^ arrays, so training can choose the resolution level during execution. General-purpose engines like DuckDB and Polars can read the same files, and a single store can support both exploratory analysis and model training. For the complete 23,546-neuron male adult nerve cord (MANC) store ^18,19^, reading the vertex positions of an entire pyramid level took 0.91 ms per neuron, 22 to 197 times faster than decoding the same meshes through 3D Tiles, Neuroglancer ^20^, or source files. A 3 × 3 tile window was read nine times faster than the full level (Methods). Linking geometry to records takes one query and an in-memory join. Returning the in-window geometry of all 11,745 neurons that intersect that window, joined to their synapse-count records, took 2.1 s, compared with approximately 16 min when those neurons were fetched individually as whole meshes from Neuroglancer precomputed files. Extended Data Table 1 compares this combination of a queryable vector-geometry pyramid and a shared semantic model across 2D and 3D with existing formats.

### One ingest serves both analysis and visualization, with full-precision geometry on the analysis path

For any dataset, one spatial index, an octree for 3D geometry or a quadtree for 2D, assigns features to cells and drives every encoder. From that index, 3D data are written to OGC 3D Tiles, Neuroglancer precomputed meshes, and a columnar Parquet store, and 2D data as conventional vector tiles and Parquet. Although the 2D and 3D routes use separate generators, they share the muDM model, descriptor, and columnar schema. The cost of indexing and cell assignment is therefore incurred once per dataset instead of separately for each output format.

For analysis, geometry remains in full precision and is not quantized. Faces are stored whole, so the finest level reproduces the source mesh. For the 23,546 MANC neurons, the unique vertices and faces recovered from the Parquet store are identical to those of the source meshes (Methods). Formats intended for rendering use quantization; Meshopt can also store coordinates bit-for-bit (Methods). Levels of detail are built by quadric error simplification for 3D meshes, by deterministic subsampling that halves the detections at each coarser level for 2D points, and optionally by Douglas–Peucker simplification for 2D polygons (Extended Data Fig. 1). The deployed catalog does not simplify polygons.

### Coarser levels retain most classification accuracy at a fraction of the storage

Since every level of detail in muDM is independently addressable, a workload can measure how much geometric detail a given task needs, a question that is otherwise costly to answer. Using the MANC connectome, which contains 23,546 neuron meshes, a compact point-cloud classifier (a dynamic graph convolutional neural network, DGCNN ^21^) was trained at each pyramid level to infer neuron type from shape under a fixed input budget. Three classification tasks were evaluated: broad neuron class, subclass, and developmental hemilineage. For each, accuracy generally rose with resolution within seed-to-seed variation, so the finest level was the most informative, and the question becomes how far a task can be coarsened before accuracy declines (Fig. 2; Table 1).

**Fig. 2.**
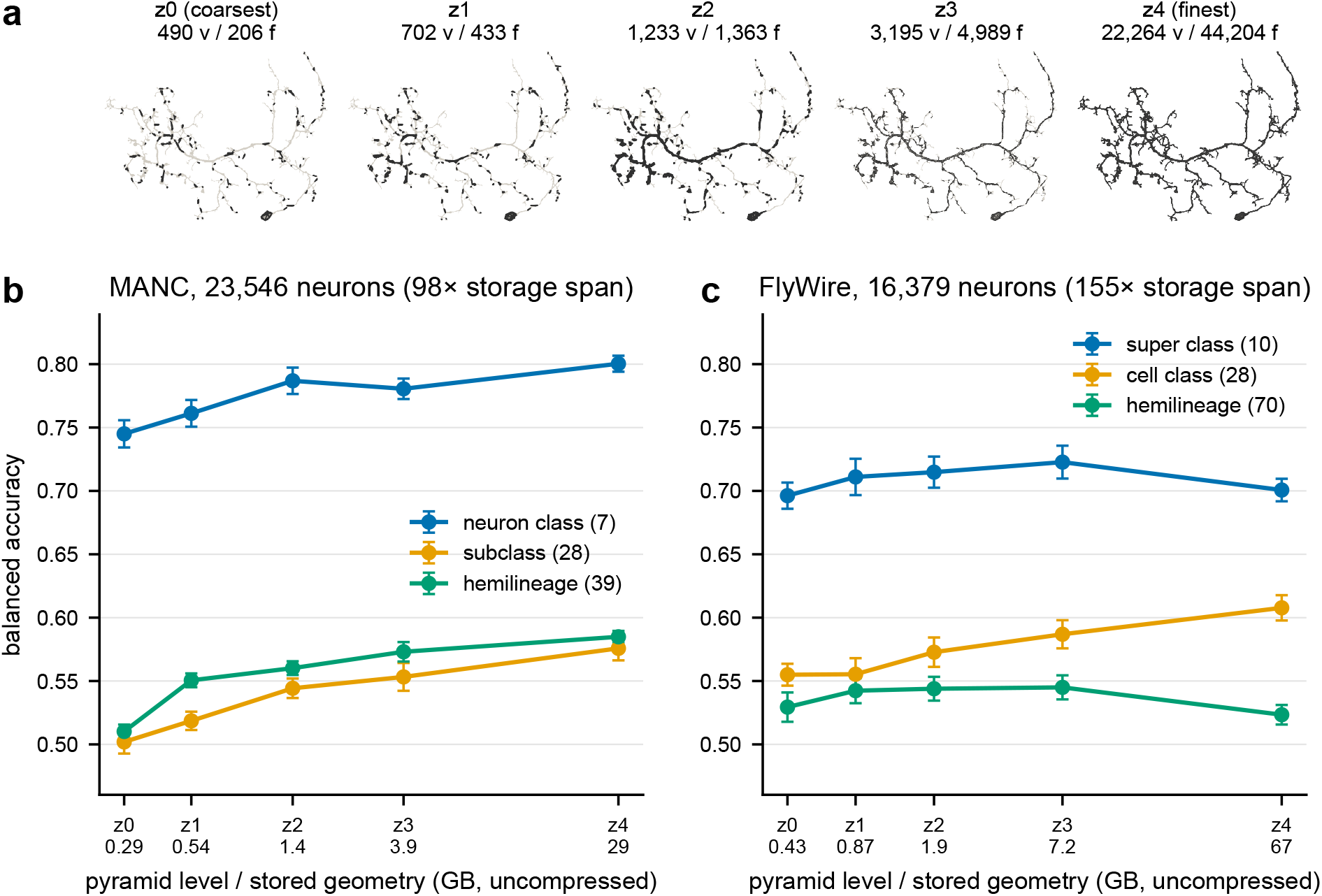
Coarser pyramid levels retain most neuron-typing accuracy at a fraction of the storage. **a**, One MANC neuron (cell type IN13A065, near the median size) at the five pyramid levels, with unique vertex (v) and face (f) counts; coarse levels consist of small, open triangle patches (Methods); the finest level is shown in light gray beneath the coarser ones. **b**, Balanced accuracy versus stored geometry per level for the MANC connectome (23,546 neurons; three tasks, DGCNN, 512-vertex budget, mean *±* s.e.m. over 15 seeds; values in Table 1). **c**, The same for a 16,379-neuron FlyWire subsample (Extended Data Table 2). Storage is the uncompressed geometry of each level on a logarithmic axis.

**Table 1.** Balanced accuracy versus storage throughout the MANC 3D-mesh pyramid (DGCNN, 512-vertex input budget, 15 seeds; balanced accuracy reported as the seed mean, s.e.m. ≤ 0.02). Storage is the uncompressed geometry of each level (98-fold span); on-disk size after ZSTD compression spans 62-fold.

| Level | Storage (GB) | On disk (GB) | Neuron class (7) | Subclass (28) | Hemilineage (39) |
| --- | --- | --- | --- | --- | --- |
| z0 (coarsest) | 0.29 | 0.16 | 0.745 | 0.502 | 0.510 |
| z1 | 0.54 | 0.29 | 0.761 | 0.519 | 0.551 |
| z2 | 1.39 | 0.72 | 0.787 | 0.544 | 0.560 |
| z3 | 3.89 | 1.93 | 0.781 | 0.553 | 0.573 |
| z4 (finest) | 28.55 | 10.14 | 0.800 | 0.576 | 0.585 |

The trade-off is favorable. Moving from the finest to the coarsest level reduces storage approximately 98-fold while balanced accuracy falls slowly. Across the three tasks, the coarsest level holds about 1% of the geometry while achieving 87 to 93% of the finest-level accuracy, with intermediate levels tracing the rest of the curve (Table 1). For the broad seven-class problem, accuracy loses only about two points through the middle of the pyramid.

Generalizability beyond MANC was tested using the same procedure on a second, independently reconstructed connectome, a class-stratified sample of 16,379 neurons from the FlyWire adult-brain connectome ^22,23^. This sample covered ten super-classes, and a leakage-safe split withheld complete cell types (Extended Data Table 2). Coarser levels again preserved most of the finest-level balanced accuracy. The coarsest level, holding roughly 0.6% of stored geometry across a 155-fold span, reached 91 to 101%, so the storage benefit was at least as favorable as for MANC. The two connectomes differed in the contribution of resolution. In FlyWire, balanced accuracy was mostly unchanged across levels for super-class and hemilineage classification, with the coarsest level indistinguishable from the finest (paired *p* ≥ 0.68); resolution improved performance only for the cell-class task. MANC improved with resolution for every task. In FlyWire, finer geometric resolution was largely unnecessary for neuron typing, which strengthens the storage economy and shows that the finest level is not always the most informative.

muDM makes this relationship inexpensive to characterize because every level is precomputed and separately addressable. Applications can evaluate candidate levels and read the selected one straight from storage, without re-tiling. The economy is in storage and serverless-read cost; forward-pass computation and memory decrease whenever the finest mesh is subsampled, with or without a pyramid, so coarsening trades accuracy for storage. Both demonstrations are 3D meshes from *Drosophila*; other species have not been tested for this task. Polygon level of detail is provided as an opt-in feature and is evaluated through geometric properties and downstream feature stability. For ∼187,000 PanNuke nuclei ^24,25^ at a level preserving 11% of vertices, per-nucleus rankings for nuclear area, Feret diameter, eccentricity, and mean intensity were maintained (Spearman *ρ* ≥ 0.99), whereas boundary-sensitive measures such as solidity and circularity were the first to degrade (Extended Data Fig. 1, Extended Data Table 3; Methods).

### Linked features improve classification beyond geometry alone

Each muDM feature keeps its measurements linked to its geometry. To quantify the benefit of this association, the neuron-typing experiment was repeated with the classifier using four synapse-count measurements per neuron in addition to the mesh: log-transformed presynaptic, postsynaptic, and total counts, plus the presynaptic fraction. These were joined to neurons through feature identifiers ^18,19,26^. Results were compared against two prespecified baselines under the same seeds and splits: the geometry-only classifier described previously, and a counts-only model (Methods). Linked counts improved balanced accuracy at every pyramid level for neuron class and subclass, and at one level for hemilineage after multiple-comparison correction (Extended Data Table 4). Gains were 0.07–0.12 for broad neuron class and 0.04–0.09 for subclass; some of these improvements may be inherent, since connectivity contributes to these class systems. Developmental hemilineage improved by as much as 0.03. Counts alone reached only 0.067 on this task, so the gain comes from combining them with geometry. In the broad classification task, the coarsest level with counts exceeded performance from geometry alone at the finest level (0.852 versus 0.800, paired *p* = 4 × 10^−3^); for subclass the difference did not reach significance (paired *p* = 0.08), and for hemilineage the finest level with geometry alone remained better. Randomly permuting counts among neurons eliminated the improvements, and the effects remained when whole cell types were withheld by the split (Methods).

### One model represents diverse spatial data at scale in a public, queryable catalog

To demonstrate that the model is not specialized for a single modality, the tooling was used on a public catalog containing more than 100 datasets across the two regimes it aims to combine. Most are 2D datasets, including multiplexed protein imaging data (CODEX, MIBI) and spatial transcriptomics datasets (Xenium ^2^, CosMx, MERSCOPE, Visium). The 3D collection comprises *Drosophila* connectome mesh reconstructions, including the FlyWire, MANC, male central nervous system (CNS), brain and nerve cord (BANC), and hemibrain connectomes, as well as whole-body human anatomical meshes from the HuBMAP Human Reference Atlas ^27^. All datasets enter the same muDM pyramid and descriptor and are delivered through one static, serverless (S3 object storage) deployment. A unified catalog interface consequently exposes both micron-scale cell boundaries and millimeter-scale neuron arbors (Extended Data Fig. 2; Supplementary Note 3). Three connectomes are served complete: the 23,546-neuron MANC connectome used for the experiments above, the FlyWire brain with all 139,244 annotated neurons ^22,23^, and the male CNS with all 176,570 neuron bodies of release v0.9 ^28^.

The catalog also serves the 134,942 BANC neurons with published meshes ^29^ and the 25,000 largest traced hemibrain bodies ^1^. All five are served as 3D Tiles and a Parquet pyramid.

The catalog, being muDM-native, columnar, and accessible through HTTP range requests, can also be used outside browser viewers. A machine-readable columnar index, queryable in place using engines such as DuckDB, together with per-dataset descriptors, enables programs to identify and retrieve only necessary datasets and resolution levels. Those descriptors also permit discovery and querying by large language model (LLM) agents, for instance via the Model Context Protocol ^15,30^.

## Discussion

muDM is designed to complement the formats already used in spatial biology. Imaging keeps its multiscale raster representations (OME-Zarr and OME-NGFF ^4^); expression matrices remain in columnar, cell-centric stores (AnnData, TileDB-SOMA ^12^); and interactive inspection continues through viewers such as Celldega ^5^, Neuroglancer ^20^, and OGC 3D Tiles clients ^9^. What these systems lack is a layer that keeps every geometric feature linked to its records and its parent structure, in both 2D and 3D and at every resolution, in a form a query engine can read directly. muDM provides this layer. It stores, selects resolutions for, and serves geometries produced by segmentation and reconstruction. It has a place complementing other tools in a processing workflow, for example, after Baysor ^31^ has assigned transcripts to cells or Bento ^32^ has analyzed their subcellular distribution, both of which operate on raw molecule tables before tiling. Table 2 outlines the muDM output appropriate to each consumer.

**Table 2.**
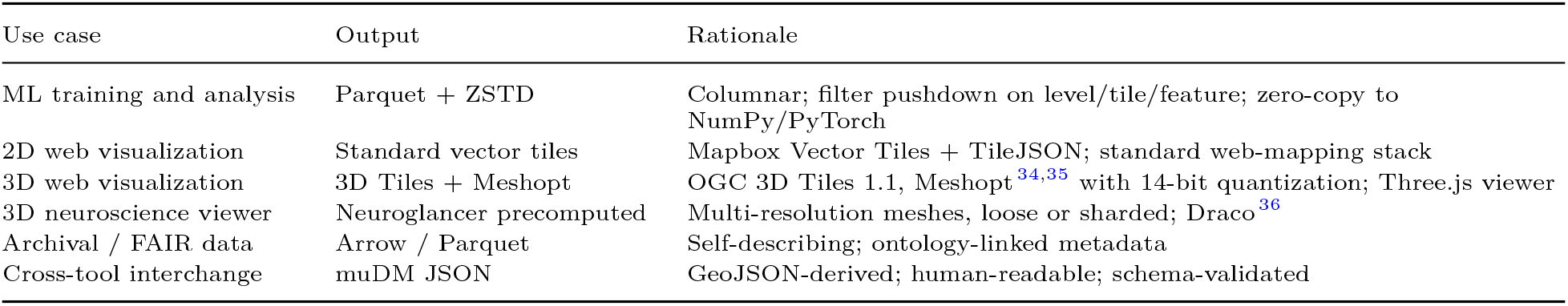
Guidance for format selection: which muDM output is suited to each use case. FAIR, findable, accessible, interoperable, and reusable.

Right-sizing provides economy in storage and reading. Required resolution depends on both the dataset and the task. For the MANC connectome, performance increased with resolution, and no task was adequately served by the coarsest level, so right-sizing there means selecting an intermediate level. By contrast, coarse geometry was generally adequate for the FlyWire brain connectome, where the lowest-resolution level matched the finest level for most tasks. The economy is observed in two connectomes, both *Drosophila* from independent reconstructions of distinct regions. Deployment currently runs on a single node, and the rendering-mesh encoders need several times the memory of the columnar path (Methods). Quadric error simplification ^33^ suits originally smoother surfaces; on the thin neurites of reconstructions it reduces each mesh chunk to a few triangles at the coarsest levels (Methods). The fusion finding has separate limitations. Its strongest improvements occur for class systems defined partly by connectivity. The hemilineage improvement, which does not have this qualification, is modest, and the demonstration covers only one organism and one linked modality. It does demonstrate that the same read path provides the model with measurements for a geometry at every resolution.

Nothing in this mechanism is tied to an organism or an assay. The same data model and pyramids could hold atlases of any tissue from any species, with every scale of a structure, and every record attached to it, a single query away.

## Code availability

Both the muDM data model and the mudm-tools tiling pipeline are released as MIT-licensed open source. The data model is available at https://github.com/NovagenResearch/mudm, while the pipeline, web viewers, and benchmark scripts are available at https://github.com/NovagenResearch/mudm-tools. A companion repository ^37^ at https://github.com/bengtl/mudm-paper-code provides the analysis code reproducing this paper’s quantitative findings, including the neuron-typing economy and fusion sweeps, the counts-only leakage probe, polygon feature-stability analysis, deposited result artifacts, and scripts for generating tables and figures. The code is MIT licensed; dataset-derived artifacts retain their upstream licenses, as described in the repository’s data-license files. The analyses used `mudm` and `mudm-tools` v0.6.4 from PyPI. The format specification is hosted at https://novagenresearch.github.io/mudm/. Release v1.0.1 of the companion repository is archived on Zenodo (https://doi.org/10.5281/zenodo.23421462).

## Data availability

A public, browsable catalog distributes the tiled datasets at https://ngrf.mudm.io. For every dataset, the catalog provides muDM tile-geometry Parquet files, ZSTD-compressed and directly readable with PyArrow, together with render-ready tiles and machine-readable descriptors. The level-of-detail experiment used the MANC *Drosophila* ventral-nerve-cord connectome ^18^, accessed through neuPrint ^26^. The catalog serves this connectome, with its Parquet pyramid listed in https://ngrf.mudm.io/tiles/manc-full/tilejson.json, and the companion repository regenerates the experimental pyramid from source. Provenance and licenses for each dataset are documented in the catalog and in Extended Data Table 5. Tiles derived from FlyWire are distributed under the CC BY-NC 4.0 license of the FlyWire public release, with attribution to the underlying source publications ^22,23,38^. The 16,379-neuron subsample used for Extended Data Table 2 came from that release. It is not redistributed; the companion repository regenerates it from source. For polygon feature-stability analysis, the PanNuke histopathology dataset ^24,25^ (CC BY-NC-SA 4.0) was ingested as muDM polygons as described in Methods. Per-neuron synapse counts for the fusion experiment were obtained from neuPrint (`manc:v1.2.1`, retrieved 11 August 2026); the resulting feature table, along with the fusion, counts-only, and leakage-probe artifacts, is deposited in the companion repository.

## Acknowledgements

The authors thank the public data providers whose datasets are included in the muDM catalog. Dataset-specific provenance and license information appear in the catalog and in Extended Data Table 5. We thank our colleagues from NGRF and Axle Informatics for their feedback during the development of muDM and during manuscript revision.

## Author contributions

B.L. developed and implemented the muDM data model and mudm-tools tiling pipeline; conducted the experiments and analyses; created the deployed catalog and web viewers; and wrote the manuscript. N.H. supervised the project, contributed to its design and evaluation, and revised the manuscript. N.S. originated the initial idea, supervised the work, contributed to design and evaluation, and revised the manuscript.

## Methods

### The muDM data model

muDM provides a spatial-annotation model derived from GeoJSON. It uses a deliberately small collection of constructs, with Pydantic v2 models ^39^ enforcing validation during construction, and is provided as the Python package `mudm`. Nine geometry classes are specified. Seven of them (Point, MultiPoint, LineString, MultiLineString, Polygon, MultiPolygon, and GeometryCollection) retain GeoJSON semantics unchanged and may include Z coordinates. Mesh geometry is represented by two 3D surface forms derived from ISO 19107: TIN (Triangulated Irregular Network), a triangular surface whose faces are closed three-vertex rings, as in GeoJSON polygons ^16^, and PolyhedralSurface, a mesh with arbitrary polygonal faces that extends TIN beyond triangular tessellations. For Parquet and binary glTF (GLB) emission, the tiling process transforms a TIN into an indexed mesh: a flat single-precision coordinate buffer accompanied by 32-bit index triples, with vertices deduplicated exactly. This representation corresponds directly to GPU vertex buffers.

Feature, inherited from GeoJSON, is the fundamental annotation object, comprising a geometry, an unrestricted properties mapping, and an optional identifier. Beyond the Feature definition, muDM adds a cross-reference identifier, a parent identifier to support hierarchical structure, and a type discriminator. Arbitrary annotation-specific information, including anatomical labels, morphometric values, provenance, and display color, resides in the properties mapping. Ontology terms, each with a URI, label, and description, are grouped into vocabularies that a feature collection carries into its JSON and GeoParquet outputs (Supplementary Note 6). Neuron or cell annotations can cite Cell Ontology terms that resolve to ontology URIs, and those references persist through tiling, serialization, and visualization. In place of GeoJSON’s WGS84 coordinate system ^16^, muDM supplies a coordinate-reference system that accommodates pixel space, physical space calibrated in micrometers or nanometers, and custom coordinates with user-defined axes and transforms. Registration of 3D volumes is supported by both an affine-transform model and a voxel coordinate system.

One shared in-memory model can be written in multiple output formats: JSON, as a GeoJSON-derived interchange dialect; Mapbox Vector Tiles for 2D viewing; glTF/GLB ^40^ with Meshopt for browser-based 3D rendering; Apache Parquet ^41^ compressed using ZSTD ^42^ for ML training and columnar analysis; and Neuroglancer precomputed meshes for volumetric visualization.

### Multi-resolution vector pyramid construction

In addition to the core data model, optimized tools are provided that transform muDM features into a multi-resolution, spatially indexed pyramid, as a Python package, `mudm-tools`. Bounding boxes determine octree-cell assignment in 3D and quadtree assignment in 2D. Every cell is keyed by its zoom level and its x and y indices, plus a depth index in 3D. Features crossing multiple cells are placed in each relevant cell. Surface-mesh faces are assigned whole to every cell they overlap, so tiling preserves geometries, and a neuron’s rows concatenate back to its source geometry with duplicated border faces (3.3% of faces and 5.9% of vertices at the finest MANC level). Polygons are clipped at 2D tile borders with the Sutherland–Hodgman algorithm ^43^, using a configurable border buffer to prevent visible seams.

Three reduction methods lower resolution as zoom becomes coarser. An edge-collapse implementation ^44^ of Quadric Error Metrics ^33^ simplifies 3D surface meshes while minimizing geometric error; because only edges are collapsed, each mesh component stays separate. At *k* levels below the finest, simplification targets 1/3^*k*^ of a neuron’s source faces; the Parquet writer simplifies each coarse-level fragment again, so the coarsest level holds under 1% of the source faces. Because connectome source meshes are already chunked into many components (a median of about 100 per MANC neuron), the coarse levels are sets of small triangle patches, roughly one per source chunk at the coarsest level, that sample the arbor’s extent. 2D polygon rings are reduced using Douglas–Peucker ^45^ toward a configurable minimum-vertex floor. The result is one simplified 2D polygon per level of detail (Extended Data Fig. 1). Point layers are thinned per feature. At *k* levels below the finest, a detection is kept if its identifier is divisible by 2^*k*^ . The rule is deterministic and preserves composition, as across three Xenium sections ^2,46,47^ (19–43 million transcripts, 541 codewords including control and blank probes), per-codeword proportions at the coarsest point level deviated from the finest by 1.0–1.5% (median) with no codeword lost, whereas per-tile counts vary binomially and 5–8% of tile–gene cells holding at least eight detections became empty (companion repository). Rust implementations, exposed through PyO3 ^48^ and parallelized with Rayon, handle clipping, decimation, transformation, and encoding.

### Storage and zero-copy access

The tiled geometries are stored using Apache Parquet ^41^, compressed with ZSTD ^42^ at level 3. Each row records tile coordinates, feature identifier, a geometry-type code, packed single-precision coordinates, packed 32-bit indices, and a tag mapping keyed by strings; the zoom level is the partition key. The 139,244-neuron FlyWire brain (728 GB of source meshes) was tiled into a five-level Parquet pyramid in 5 h 4 min with a peak resident size of 33 GB on the benchmark host. The rendering-mesh encoders hold more state, peaking at 141 GB (3D Tiles) and 200 GB (Neuroglancer) for the 23,546-neuron MANC set. Files are partitioned into one directory per zoom level (Hive layout), and filter pushdown on zoom, tile coordinates, and geometry type reads only the row groups a task needs, for example coarse whole-arbor meshes at zoom 0 or fine local meshes at zoom 3. The appropriate level is selected by comparing accuracy among levels. Packed coordinate and index byte columns can be exposed zero-copy through PyArrow as NumPy arrays and PyTorch ^17^ tensors, eliminating deserialization for a single row; assembling a neuron from several rows concatenates them. One ingest produces 3D Tiles, Parquet, and Neuroglancer precomputed meshes from the same octree. Fidelity of the finest level was checked directly by hashing vertex coordinates and face vertex-triples and comparing the sets of unique vertices and unique faces recovered from the Parquet store with those of the source OBJ files for every MANC neuron (result deposited in the companion repository).

### Mesh codecs

For browser display, cell-assigned mesh data are written as glTF 2.0 GLB files in an OGC 3D Tiles hierarchy and use the Meshopt GLB extension ^34^. Meshopt can retain coordinates bit-for-bit, but the deployed catalog applies visually lossless 14-bit positional quantization ^49^ to reduce size. Tilesets pass the OGC 3D Tiles validator (Supplementary Note 1).

Neuroglancer precomputed meshes are written as per-neuron files, the layout distributed and benchmarked here, or in the sharded format ^50^, and are compressed with Draco ^36^. Positions are pre-quantized to an integer grid and Draco adds no further quantization, so decoding returns the gridded positions exactly (Supplementary Note 5). The sharded output conforms to the specification and renders in the official Neuroglancer client (Supplementary Note 1).

### muDM neuron-typing protocol

Neuron surface meshes from the MANC *Drosophila* ventral-nerve-cord connectome ^18,19^ formed the basis of the right-sizing study. The meshes are level 2 (of 0–3, 0 finest) of the published MANC multi-resolution meshes, with labels from neuPrint ^26^, and cover 23,546 traced, typed neurons; FlyWire meshes were taken at level 1. Every mesh was placed into a five-level octree pyramid, with zoom level 0 as the coarsest and zoom level 4 as the finest. Three neuPrint-derived labels supplied classification tasks at distinct granularities: neuron class with 7 classes, subclass with 28, and developmental hemilineage with 39. For each target, classes containing fewer than 20 members were removed. After this filter and the omission of neurons without an instance annotation (below), 21,946, 20,616, and 14,402 neurons remained for neuron class, subclass, and hemilineage, split 70/15/15 into training, validation, and test sets for each seed (mean test sets of 3,266, 3,095, and 2,162 neurons). Cell type (117 classes) was excluded because its classes coincide with the homolog groups used to prevent leakage, so no leakage-safe split exists for that task; a per-neuron split, which cannot exclude bilateral twins, is reported for reference in Supplementary Note 2.

To avoid leakage between bilateral homologs, training, validation, and test partitions were grouped according to the instance annotation after removing the left/right side suffix. Mirror-symmetric pairs therefore always occupy one split. Neurons without an instance annotation were omitted, as singleton treatment could separate a pair across partitions. Partitions were stratified by class and re-sampled for each seed. Serial homologs can still share geometry because they have the same cell type across neuromeres while possessing distinct instance annotations and therefore are not grouped. Across 15 seeds, no split group crossed the train/test boundary. However, 38 to 42% of test neurons shared their cell type with a training neuron from another group. Because all resolution levels use the same fixed split, such exposure is identical at every level. Repeating all three sweeps with grouping by cell type, holding out entire types so that neither bilateral nor serial homologs occur in both train and test, gave the same trade-off. This split is reused for the fusion controls. The trade-off was also preserved with a PointNet classifier ^51^ under the identical protocol (matching or exceeding DGCNN retention), at input budgets of 2,048 and 4,096 sampled vertices, and with a 30-epoch recipe using validation-based epoch selection, which raised absolute accuracy at every level without reducing coarsest-level retention at 512 vertices (Supplementary Note 2). Because absolute accuracy decreased at larger budgets with both training lengths, and longer training left the trade-off unchanged, Table 1 uses the 512-vertex, eight-epoch recipe.

The classifier is a small DGCNN ^21^. It consists of EdgeConv layers operating on a *k*-nearest-neighbor graph (*k* = 20, capped at the smallest point count in each batch), followed by global max pooling and a two-layer classification head. At each level, each neuron contributed no more than *B* = 512 sampled vertices. Neurons with fewer than *B* vertices used all of them, and batch padding was excluded from neighbor search and pooling. Input points were drawn uniformly at random, without replacement, from the concatenated rows of the level being evaluated, without removing border duplicates (5.9% of vertices at the finest level); the finest level, whose unique vertices and faces match the source mesh, served as the reference. Training used Adam ^52^ with learning rate 10^−3^, batch size 64, and eight epochs, across 15 seeds, on an 80-core Linux system equipped with two NVIDIA A4500 GPUs. Because class frequencies are imbalanced, balanced accuracy, defined as mean per-class recall, is reported as mean ± s.e.m. across seeds. The companion repository contains the per-seed values underlying every reported mean.

### Economy and right-sizing readout

Storage for each level was obtained directly from Parquet column metadata, without decompression. At every zoom level, the uncompressed sizes of all columns were summed over row groups. Table 1 lists the uncompressed geometry and the on-disk ZSTD-compressed size, the quantity moved over a network, for each level. At every pyramid level, balanced accuracy was assessed against the finest-level reference and reported by task and level as the seed mean with standard error (Table 1). Levels were compared with paired *t*-tests ^53^ over the 15 seeds. The benefit of loading a coarser level is expressed as the ratio of the finest level’s stored geometry to its own.

### Read-path benchmark

The read-path measurements reported in Results used the complete MANC dataset of 23,546 neurons. On the benchmark host’s local disk, every output format was generated from the same shared tile index by the released mudm-tools v0.6.4 pipeline. The columnar arm scanned the feature-identifier and vertex-position columns of the finest level once; the per-item arms decoded complete meshes. Per-item representations (3D Tiles, Neuroglancer precomputed, and source OBJ) were benchmarked on a 2,000-neuron subset and scaled linearly to a full level; the one-time 3D Tiles scan that maps neurons to tiles was counted once. For individual-feature retrieval, 50 identifiers were each fetched five times. The feature-to-record join executed a single filter-pushdown query over a 3 × 3 tile window at the finest level, then joined every returned feature in process to its per-neuron synapse-count record. The render-format comparator received the feature list for that window without cost, retrieved each neuron’s complete finest-level mesh separately, and was timed for 200 neurons before linear extrapolation. Per-object access favors Neuroglancer: with a warm page cache, its per-neuron files return one feature in a median of 10 ms, compared with 83 ms for a level re-sorted by feature identifier and 330 ms for the default tile-sorted layout. All artifacts are available in the companion repository.

### Multimodal fusion protocol

For the fusion study, the neuron-typing procedure was rerun with the DGCNN receiving four scalar values per neuron in addition to the point cloud: log-transformed presynaptic, postsynaptic, and total synapse counts, together with presynaptic fraction. These values came from MANC neuPrint annotations ^18,26^. In the deployed catalog, the counts are stored as feature tags; in the training harness, they were provided through a per-neuron table indexed by the same body identifier. The four values are appended to the pooled global representation before the classifier head. Two comparators used the same partitions, seeds, epoch counts, and input budgets: the geometry-only model from Table 1 and a counts-only multilayer perceptron using those four scalars. Before the main experiment, a leakage probe deposited with the companion code estimated the predictive signal available from counts alone by inferring every task label from counts under the same grouped split. For each level, paired *t*-tests ^53^ over the 15 shared seeds were performed and Holm-corrected ^54^ across the five levels within each task. All neuron-class and subclass improvements, plus the level 3 hemilineage improvement, remained significant after correction; the coarsest hemilineage improvement narrowly missed significance after adjustment (*p* = 0.052).

Two controls assess the comparison. First, shuffling count features among neurons using one fixed permutation removed the benefit. The maximum absolute change across all 15 task–level cells was 0.013, and no cell survived Holm correction. Improvement therefore requires that counts be associated with the correct neurons, not merely that additional model input capacity be present. Second, applying the fusion experiment to the cell-type-held-out partition used for the economy robustness test preserved its effects. Neuron-class gains ranged from 0.08 to 0.12 and subclass gains from 0.05 to 0.09 at every level, all with *p* < 1.8 × 10^−4^. Hemilineage gains remained positive but small across all five levels and were Holm-significant at three.

### Cross-format round-trip and 2D breadth

Round-trip equivalence was evaluated by serializing identical features through JSON, vector tiles, glTF/GLB, and Parquet, recovering their geometry, and verifying geometric equivalence. Spatial-transcriptomics and histopathology datasets exercised 2D coverage. A 10x Xenium formalin-fixed paraffin-embedded (FFPE) bundle was imported with cell and nucleus outlines as 2D polygons, each tagged with its cell identifier, which keys the cell’s expression vector in a Parquet store served beside the tiles. Each transcript detection is stored as one point feature tagged with its gene name, from three levels below the finest zoom to the finest. Since Xenium lacks a native cell-type ontology, its cells are represented primarily through expression. The breast-cancer Rep1 dataset ^2^ contained 167,780 cells and a 313-gene panel. The same ingest produces Mapbox Vector Tiles for 2D visualization, an image-channel raster pyramid, and Parquet for ML. PanNuke nucleus polygons ^24^ provided the histopathology example and support the 2D polygon level-of-detail analysis in Extended Data Fig. 1.

### Feature robustness across polygon levels

Extended Data Table 3’s feature-stability analysis included 7,556 PanNuke patches converted to muDM polygons (∼187,000 nuclei). Douglas–Peucker simplification toward the pipeline’s minimum-vertex floor was applied to every nucleus polygon at tolerances of *τ* ∈ {0.5, 1, 2, 4, 8} pixels. Each result was then rasterized into a 16-bit instance mask on the original 256 × 256 pixel grid. A nucleus received the same label at each level, permitting per-nucleus rows to be joined across levels. Nyxus v0.11.0 ^55^ calculated 144 numeric morphology and intensity features per nucleus at every level against the original grayscale intensity image. For each feature at each level, agreement with full detail is reported using the median absolute relative deviation over nuclei and the Spearman rank correlation ^53^ of per-nucleus measurements. Nuclei were linked between levels by their (patch, label) pair, and features with zero variance were omitted from correlation analysis. Because both levels in a comparison share the rasterization stage, most discretization errors disappear.

