## Supplementary Information for "muDM: A Queryable Multi-Resolution Data Model for Vector Geometry in Spatial Bioimaging"

Superscript numbers refer to the reference list of the main article.

#### Supplementary Note 1: Mesh codec implementation and conformance

**3D Tiles and GLB.** Meshopt[34,35] is a byte-oriented codec employing delta, zigzag, and varint coding. Tilesets were checked with 3d-tiles-validator 0.6.1 and loaded in CesiumJS 1.121 and Three.js.

**Neuroglancer precomputed meshes.** Meshes are generated either as loose per-neuron files or in the sharded form. To assign mesh labels to shards and minishards, the sharded writer uses the same MurmurHash3 variant used by CloudVolume and Neuroglancer. The sharded output was tested end-to-end in the live Neuroglancer viewer: a sharded store produced by the mudm-tools pipeline was served through HTTP range requests, then loaded and rendered in the official Neuroglancer 2.41.2 client.

#### Supplementary Note 2: Robustness of the level-of-detail trade-off

All variants follow the MANC neuron-typing protocol (Methods) except for the factor varied. Retention is coarsest-level balanced accuracy expressed as a percentage of finest-level balanced accuracy for the same configuration. Per-seed values for every variant are deposited in the companion repository.

- **Cell-type grouping.** With entire cell types held out of training, coarsest-level retention remained within a few points of that observed with instance grouping.
- **Classifier.** Under the identical protocol, PointNet[51] retained 94, 93, and 99% of its own finest-level balanced accuracy at the coarsest level for neuron class, subclass, and hemilineage, matching or exceeding the DGCNN fractions (93, 87, and 87%; main-text Table 1).
- **Input budget.** Coarsest-level retention for neuron class, hemilineage, and FlyWire super-class was 93, 87, and 99% at 512 sampled vertices and 98, 97, and 99% at 4,096. Absolute accuracy decreased at larger budgets with eight-epoch training and, in a three-seed neuron-class run, with 30-epoch training.
- **Training length.** Training for 30 epochs with validation-based epoch selection raised absolute accuracy at every level (neuron class: from 0.75 to 0.87 at the coarsest level and from 0.80 to 0.90 at the finest), with coarsest-level retention of 91 to 96% across the three MANC tasks at 512 vertices and 93% at 2,048

in a three-seed neuron-class run. For hemilineage, the three intermediate levels then came within 2.3% of the finest level.

- **Cell type.** Cell type (117 classes) has no leakage-safe split (Methods). A per-neuron split, which cannot exclude bilateral twins, gave 0.40 balanced accuracy at the finest level with 76% retained at the coarsest.

### Supplementary Note 3: Web visualization

A Three.js web viewer displays 3D Tiles output using level-of-detail traversal, frustum culling, feature picking, and a bounded GPU cache (Extended Data Fig. 2). On each frame, the viewer culls tiles outside the viewing frustum and, for each selected feature, selects the coarsest level whose screen-space error against the tile bounding volumes falls below a tuned threshold; tiles already shown stay visible until the new level loads. Only tiles containing selected features are loaded. A configurable GPU-memory limit constrains resident GLB data and triggers least-recently-used tile eviction when memory pressure occurs. Clicking a feature reveals properties held in glTF node metadata, including cell type, instance, body identifier, synapse counts, and display color.

### Supplementary Note 4: Structural analysis of columnar versus scene-graph encoding

This section makes the structural case for the muDM columnar Parquet schema as an ML-training representation relative to scene-graph glTF, the standard web container for 3D. The comparison is structural, apart from one timing measurement on shared data, and covers the correspondence between each layout and common training-loop tasks: building tensors, looking up categories, and parallel loading.

**Flat arrays to direct tensor mapping.** In muDM's tile-geometry schema, mesh triangle indices and vertex coordinates are stored as contiguous packed-binary columns of single-precision floating-point and integer values. With a single zero-copy buffer-to-array call, a consumer can map such a column into a NumPy or PyTorch array without reshaping nested structures. In glTF, equivalent data sit behind a chain of indirections (node, mesh, primitive, accessor, buffer view) before the binary buffer, and a JSON manifest must be parsed for every tile to resolve this chain. The format-level effect appears on the 23,546-neuron MANC dataset, with every format encoded from a shared tile index. Reading the vertex positions of an entire pyramid level takes 0.91 ms per neuron from the columnar store, compared with 20.4 ms per neuron to decode the meshes via the 3D Tiles scene graph and 70.0 ms through Neuroglancer per-neuron files (Methods; companion repository). At the schema level, columnar organization eliminates that chain of indirections.

**Dictionary indices to category lookup.** Tag keys and values are dictionary-encoded within each Parquet column chunk, so each unique tag string is stored once, with integer references elsewhere. Readers can request these columns as Arrow dictionary arrays and build a category index from the dictionaries instead of parsing every feature's strings. In glTF, equivalent metadata occurs as JSON key-value strings per feature within each glTF node's extras object, and consumers must parse the strings and build their own indices.

**Uniform schema to parallel data loading.** A single columnar schema applies to every tile, so one parsing function handles any tile in a PyTorch data loader and workers can partition work without content-dependent branches. By design, OGC 3D Tiles can be heterogeneous: a tileset may contain glTF, Batched 3D Models, Point Clouds, and Instanced 3D Models, so a general reader requires polymorphic dispatch. The stable columnar schema eliminates this branch.

**A worked query.** Because the store is a Hive-partitioned Parquet directory, any Parquet-native engine can read it without muDM-specific code. The DuckDB query below retrieves geometries, identifiers, and tags for all features in a 3 × 3 tile window at pyramid level 3. Its zoom and tile-coordinate conditions are pushed down for row-group and partition pruning:

```
SELECT feature_id, positions, tags
FROM read_parquet('manc-full/zoom=*/part_*.mu.parquet',
                  hive_partitioning = true)
WHERE zoom = 3
  AND tile_x BETWEEN 3 AND 5
  AND tile_y BETWEEN 4 AND 6;
```

The tags field is the feature-level key-value map containing identities and measurements. Query predicates target stored fields (zoom, tile coordinates, feature identifiers, and tag values), while spatial tests finer than the precomputed tile grid are performed by the consumer on the returned geometry.

#### Supplementary Note 5: Relationship to Neuroglancer precomputed meshes

For neuroscience viewers, Neuroglancer's multi-resolution precomputed mesh representation is the community-standard way to serve triangle meshes, and muDM produces it alongside OGC 3D Tiles as a rendering output. The two rendering codecs differ: GLB and 3D Tiles rely on Meshopt, whereas Neuroglancer uses Draco.

**Draco path.** Under normal use, Draco compression quantizes vertex positions and applies prediction coding. For Neuroglancer precomputed meshes, muDM first places coordinates on an explicit integer grid and then sets Draco's coordinate-wise quantization stage to identity, exercising the Draco bitstream without introducing further loss. DracoPy/libdraco decode verification returns integer coordinates with zero quantization error and matching face counts, so Draco adds no loss beyond the integer grid.

#### Supplementary Note 6: muDM specification summary

This section consolidates specification-level information described in Methods.

**Ontology model types.** Controlled vocabularies in muDM are represented by two Pydantic model types:

```
class OntologyTerm(BaseModel):
    uri: str                          # e.g., "http://purl.obolibrary.org/obo/CL_0000540"
    label: str | None = None          # e.g., "neuron"
    description: str | None = None    # e.g., "a nervous-system cell that conducts nerve impulses"

class Vocabulary(BaseModel):
    namespace: str | None = None      # e.g., "http://purl.obolibrary.org/obo/"
    description: str | None = None
    terms: dict[str, OntologyTerm]
```

An optional vocabularies field is present on both features and feature collections, mapping property names onto Vocabulary objects. Serialization preserves vocabularies in muDM JSON, as JSON, and, for feature collections, in Arrow/GeoParquet, as schema metadata (key `mudm:vocabularies`).

**Linking fields.** Two optional fields link a feature beyond its own properties. The parent identifier (`parentId`) names the feature that contains it, expressing hierarchy such as a nucleus within a cell or a cell within a tissue region. The reference field (`ref`) holds a URI or record identifier for the external resource that carries the feature's data, for example an array store or the expression matrix containing a cell's readout. Both are foreign members from a pure-GeoJSON perspective and are ignored by GeoJSON-only consumers, so a muDM document that uses only the seven GeoJSON geometry types remains valid GeoJSON.

**Property dictionary examples.** The open properties dictionary contains metadata for individual annotations, such as cell-type classification (`{"cell_type": "Kenyon cell", "class": "intrinsic neuron"}`), morphometric measurements (`{"area_um2": 125.3, "volume_um3": 1250.0}`), provenance (`{"annotator": "cellpose_v3"}`), and display color (`{"color": "#3288BD"}`).

#### Extended Data Table 1: Related-work comparison

Extended Data Table 1 compares muDM with the formats currently used by spatial-bioimaging analysts along the dimensions that muDM brings together. No existing format supplies the full set for vector geometry. muDM is the only entry with a queryable, multi-resolution vector pyramid across 2D and 3D, and it keeps each feature's semantics and linked records attached at every level.

TileDB-SOMA is the nearest existing approach. Its SOMA spatial extension combines columnar storage, vector geometry (polygons and point clouds), and a resolution pyramid, but the pyramid is limited to raster images, and polygon and point tables remain single-resolution. Current spatial-omics stores, including SpatialData and Celldega's visualization-specific DegaFiles, retain vector features at only one resolution. Celldega's semantic zoom changes which entity type a viewer displays at each scale and leaves stored geometry resolution unchanged. Its Parquet tiles can be read by engines, but they are organized for viewport streaming,

**Extended Data Table 1** Comparison of spatial-data formats. Vec. geom.: native vector-geometry type (2D: 2D only; render: rendering input only). Vec. LOD: a vector-geometry pyramid with multiple levels of detail; raster denotes formats whose pyramid applies only to raster imagery. Queryable: columnar/SQL filter pushdown. Serverless: direct object-storage access through HTTP range reads. Formats are assessed against their core specifications as of August 2026; neither third-party extensions nor tool-specific overview conventions are included. Semantics: how feature meaning is carried (generic: application-defined properties without a shared vocabulary).

| Format | Vec. geom. | Vec. LOD | Queryable | 3D mesh | Serverless | Semantics |
|---|---|---|---|---|---|---|
| **muDM** | yes | yes | yes | yes | yes | ontology |
| SpatialData | 2D | raster | partial | no | partial | AnnData |
| OME-NGFF / OME-Zarr | no | raster | no | no | yes | image |
| TileDB-SOMA / Census | yes | raster | yes | no | yes | matrix |
| Vitessce | render | no | no | no | yes | inputs |
| Celldega (DegaFiles) | 2D | raster | partial | no | yes | viewer |
| spora | no | raster | partial | no | partial | markers |
| spatialproteomics | no | no | no | no | partial | labels |
| GeoParquet 1.1 | yes | no | yes | limited | yes | generic |
| PMTiles / MVT | 2D | yes | no | no | yes | generic |
| Neuroglancer / 3D Tiles | render | yes | no | yes | yes | generic |

and query pushdown was not a design goal. The same absence occurs in datasets expressly packaged for machine learning. The spatial-proteomics release spora, which underlies the Virtual Tissues foundation model, tiles images into pyramids but represents cell segmentation solely as full-resolution integer-labeled raster masks with no vector form, so its columnar marker tables contain no geometry. The spatialproteomics toolbox likewise maintains synchronized single-resolution arrays for images, integer label masks, and per-cell tables, and defines neither a geometry layer nor a pyramid.

### Extended Data Table 2: FlyWire second-connectome economy

**Extended Data Table 2** Second-connectome economy in FlyWire: storage against balanced accuracy throughout the FlyWire 3D-mesh pyramid (DGCNN, 512-vertex budget, 15 seeds; balanced accuracy averaged across seeds). The class-stratified sample contains 16,379 neurons, and entire cell types are held out between training and test sets (leakage-safe). Stored geometry differs by approximately 155-fold between the coarsest and finest levels (119-fold on disk).

| Level | Storage (GB) | On disk (GB) | Super class (10) | Cell class (28) | Hemilineage (70) |
|---|---|---|---|---|---|
| z0 (coarsest) | 0.43 | 0.25 | 0.696 | 0.555 | 0.529 |
| z1 | 0.87 | 0.47 | 0.711 | 0.555 | 0.542 |
| z2 | 1.93 | 1.01 | 0.715 | 0.573 | 0.544 |
| z3 | 7.16 | 3.66 | 0.723 | 0.587 | 0.545 |
| z4 (finest) | 66.97 | 30.03 | 0.701 | 0.608 | 0.523 |

### Extended Data Table 3: Feature robustness across 2D polygon levels

Extended Data Fig. 1 describes polygon level of detail in geometric terms, whereas Extended Data Table 3 and its continuation evaluate whether downstream quantitative features remain intact (Methods). The table tracks widely used nuclear morphometry and intensity measures, which fall into three regimes. Robust features include size, ellipse shape, extremal diameters, and intensity aggregates: maximum Feret diameter, eccentricity, aspect ratio, and mean intensity retain per-nucleus ordering at Spearman $\rho \geq 0.97$ through $\tau = 2$ px. Area-coupled quantities, including area and integrated intensity, decline systematically because simplification removes corners inward. Their absolute error reaches tens of percent at coarse levels. Because the bias is nearly uniform across nuclei, ranks are retained ($\rho \approx 0.98$ even at $\tau = 8$ px), so comparative analyses remain viable while calibrated magnitude measurements do not. Features reflecting boundary detail fail earliest and quantify what simplification removes. Solidity and convex-hull area already lose rank fidelity at $\tau = 1$ px ($\rho \approx 0.76$–$0.80$), followed closely by extent and circularity.

**Extended Data Table 3** Robustness of individual nuclear intensity and morphometry features over polygon levels of detail (PanNuke; ~187,000 nuclei per level). Dev., median absolute relative deviation from full detail ($\tau = 0$); $\rho$, Spearman correlation of per-nucleus ranks. Rows follow the three observed regimes. Across the 140 derived features with defined values, the median deviation at $\tau = 1$ px is 1.2%, with median $\rho = 0.99$; the companion repository provides the complete feature-by-feature table.

| Feature | $\tau = 1$ px dev. | $\rho$ | $\tau = 2$ px dev. | $\rho$ | $\tau = 8$ px dev. | $\rho$ |
|---|---|---|---|---|---|---|
| *Size, ellipse shape, and extremal diameters (robust)* | | | | | | |
| `STAT_FERET_DIAM_MAX` | 0.000 | 0.999 | 0.000 | 0.998 | 0.031 | 0.991 |
| `ASPECT_RATIO` | 0.000 | 0.990 | 0.000 | 0.978 | 0.000 | 0.863 |
| `ECCENTRICITY` | 0.014 | 0.992 | 0.020 | 0.982 | 0.054 | 0.872 |
| `ELONGATION` | 0.021 | 0.992 | 0.031 | 0.982 | 0.089 | 0.872 |
| `ROUNDNESS` | 0.021 | 0.992 | 0.033 | 0.979 | 0.153 | 0.837 |
| `COMPACTNESS` | 0.029 | 0.999 | 0.054 | 0.997 | 0.325 | 0.961 |
| *Intensity aggregates (robust; region membership shifts slightly)* | | | | | | |
| `MEAN` | 0.010 | 0.999 | 0.021 | 0.996 | 0.059 | 0.979 |
| `MEDIAN` | 0.009 | 0.999 | 0.017 | 0.996 | 0.049 | 0.981 |
| `STANDARD_DEVIATION` | 0.028 | 0.990 | 0.057 | 0.969 | 0.164 | 0.848 |
| `ENTROPY` | 0.008 | 0.962 | 0.014 | 0.921 | 0.042 | 0.719 |
| *Area-coupled magnitudes (systematic shrinkage; rank-preserving)* | | | | | | |
| `AREA_PIXELS_COUNT` | 0.042 | 1.000 | 0.084 | 0.999 | 0.347 | 0.977 |
| `INTEGRATED_INTENSITY` | 0.054 | 0.999 | 0.107 | 0.998 | 0.403 | 0.976 |
| *Boundary detail (degrades first)* | | | | | | |
| `PERIMETER` | 0.009 | 0.945 | 0.019 | 0.926 | 0.197 | 0.801 |
| `CIRCULARITY` | 0.014 | 0.843 | 0.029 | 0.764 | 0.338 | 0.351 |
| `EXTENT` | 0.038 | 0.907 | 0.075 | 0.761 | 0.315 | 0.412 |
| `CONVEX_HULL_AREA` | 0.117 | 0.800 | 0.327 | 0.707 | 0.597 | 0.482 |
| `SOLIDITY` | 0.130 | 0.759 | 0.327 | 0.649 | 0.603 | 0.475 |

A small group of measures is intrinsically vertex-count-sensitive and should not be calculated from simplified levels (Extended Data Table 3, continued panel). At $\tau = 1$ px, perimeter fractal dimension plus the minimum and mode statistics of the Martin and Nassenstein diameters have already lost most ranking information; the Feret and Martin means remain robust, whereas the Nassenstein mean falls to $\rho = 0.83$.

**Extended Data Table 3** (continued) Features sensitive to vertex count that should be avoided at simplified levels: the eight measures having the poorest rank stability at $\tau = 2$ px. Dev., median absolute relative deviation from full detail; $\rho$, Spearman correlation across nuclei. Every remaining feature with defined correlations has $\rho \geq 0.62$ at $\tau = 2$ px.

| Feature | $\tau = 1$ px dev. | $\rho$ | $\tau = 2$ px dev. | $\rho$ |
|---|---|---|---|---|
| `STAT_NASSENSTEIN_DIAM_MIN` | 1.00 | 0.06 | 1.00 | 0.05 |
| `FRACT_DIM_PERIMETER` | 0.63 | 0.27 | 0.91 | 0.07 |
| `STAT_NASSENSTEIN_DIAM_MODE` | 1.00 | 0.16 | 1.00 | 0.10 |
| `STAT_MARTIN_DIAM_MIN` | 0.50 | 0.28 | 0.50 | 0.16 |
| `ALLCHORDS_MAX_ANG` | 0.33 | 0.32 | 0.50 | 0.24 |
| `ALLCHORDS_MIN_ANG` | 0.33 | 0.32 | 0.50 | 0.24 |
| `STAT_MARTIN_DIAM_MODE` | 0.40 | 0.51 | 0.50 | 0.47 |
| `ALLCHORDS_MIN` | 0.00 | 0.67 | 0.00 | 0.58 |

### Extended Data Table 5: Companion data catalog

The muDM pipeline has processed more than 100 public datasets from end to end (`mudm-tools` v0.6.3; the deployed catalog predates the v0.6.4 version used in the paper experiments) and serves them through a

**Extended Data Table 4** Geometry fused with linked synapse counts: balanced accuracy for each task and pyramid level (DGCNN, 512-vertex budget, average of 15 seeds). Δ, fusion less geometry-only; $p$, paired $t$-test over the 15 common seeds (uncorrected). After Holm correction[54] across the five levels in each task, all neuron-class and subclass improvements remain significant, as does the hemilineage gain at z3; the z0 hemilineage improvement narrowly fails after adjustment ($p = 0.052$). Counts-only does not depend on level, so its per-level values remain flat within noise.

| Task (classes) | Level | Geometry | Counts | Fusion | Δ | $p$ |
|---|---|---|---|---|---|---|
| neuron class (7) | z0 | 0.745 | 0.499 | 0.852 | +0.107 | $6.6 \times 10^{-8}$ |
| | z1 | 0.761 | 0.496 | 0.882 | +0.120 | $7.4 \times 10^{-7}$ |
| | z2 | 0.787 | 0.503 | 0.896 | +0.109 | $8.7 \times 10^{-6}$ |
| | z3 | 0.781 | 0.503 | 0.899 | +0.119 | $1.6 \times 10^{-9}$ |
| | z4 | 0.800 | 0.499 | 0.874 | +0.074 | $7.9 \times 10^{-6}$ |
| subclass (28) | z0 | 0.502 | 0.159 | 0.592 | +0.091 | $2.0 \times 10^{-7}$ |
| | z1 | 0.519 | 0.163 | 0.584 | +0.065 | $7.4 \times 10^{-5}$ |
| | z2 | 0.544 | 0.160 | 0.603 | +0.059 | $1.1 \times 10^{-5}$ |
| | z3 | 0.553 | 0.162 | 0.612 | +0.059 | $3.0 \times 10^{-6}$ |
| | z4 | 0.576 | 0.165 | 0.615 | +0.039 | $2.4 \times 10^{-3}$ |
| hemilineage (39) | z0 | 0.510 | 0.067 | 0.530 | +0.020 | 0.013 |
| | z1 | 0.551 | 0.067 | 0.562 | +0.012 | 0.025 |
| | z2 | 0.560 | 0.067 | 0.573 | +0.013 | 0.028 |
| | z3 | 0.573 | 0.065 | 0.603 | +0.030 | $4.8 \times 10^{-4}$ |
| | z4 | 0.585 | 0.067 | 0.593 | +0.009 | 0.19 |

public data catalog at https://ngrf.mudm.io. The catalog includes both spatial dimensionalities and multiple imaging modalities. The selection in Extended Data Table 5 includes four 2D Xenium spatial-transcriptomics samples from four human tissues, one Xenium mouse sample retained to demonstrate format and chemistry, the three *Drosophila* connectomes served complete (MANC, FlyWire, male CNS), the 134,942 BANC neurons with published meshes, the 25,000 largest traced hemibrain bodies, and the two whole-body 3D anatomical atlases of the HuBMAP Human Reference Atlas.

Catalog outputs consist of muDM tile-geometry Parquet files (ZSTD-compressed; not GeoParquet), plus their associated tile pyramids. Each dataset includes upstream provenance and the generation command used to create it, so the published artifacts can be regenerated from original sources. Bundled Leaflet viewers explore 2D datasets, while bundled Three.js viewers explore 3D datasets.

**Extended Data Table 5** A selection from the companion catalog of over 100 public datasets processed by the muDM pipeline. "Scale (served)" gives the content of each served artifact after ingestion and tiling. Tx, transcripts.

| Dataset | Modality | Organism / tissue | Scale (served) | Source / provenance, license |
|---|---|---|---|---|
| Xenium human breast | 2D cell + nucleus polygons, transcript points, DAPI | Human breast (FFPE) | 167,780 cells; 42.6M tx; 280-gene panel + 33-gene add-on | 10x Genomics[2]; CC BY 4.0 |
| Xenium human ovary | 2D cells + transcripts | Human ovary | 247,636 cells; 19.1M tx; 380-gene Immuno-Oncology panel + 100-gene add-on | 10x Genomics[47]; CC BY 4.0 |
| Xenium human lung | 2D cells + transcripts | Human lung | 150,365 cells; 30.6M tx; 377-gene Multi-Tissue & Cancer panel | 10x Genomics[46]; CC BY 4.0 |
| Xenium human kidney | 2D cells + transcripts | Human kidney | 465,534 cells; 77.1M tx; Multi-Tissue & Cancer panel + protein add-on | 10x Genomics[56]; CC BY 4.0 |
| Xenium mouse ileum *(format / chemistry demonstration)* | 2D cells + transcripts | Mouse ileum | 23 cells (10x Prime 5K format-demo subset); ∼5,000-gene panel | 10x Genomics[57]; CC BY 4.0 |
| MANC | 3D TIN meshes; 3D Tiles, Parquet | *Drosophila* ventral nerve cord (male) | 23,546 neuron surfaces (complete) | Janelia FlyEM, MANC v1.2.1[18,19]; CC BY 4.0 |
| FlyWire | 3D TIN meshes; 3D Tiles, Parquet | *Drosophila* brain (whole) | 139,244 neuron surfaces (all annotated) | FlyWire 783 public release[22,23,38]; CC BY-NC 4.0 |
| Male CNS | 3D TIN meshes; 3D Tiles, Parquet | *Drosophila* whole CNS (male) | 176,570 neuron surfaces (all neuron bodies) | Janelia FlyEM, MaleCNS v0.9[28]; CC BY 4.0 |
| BANC | 3D TIN meshes; 3D Tiles, Parquet | *Drosophila* whole CNS (female) | 134,942 neuron surfaces (all with published meshes) | BANC-FlyWire Consortium, v888 public snapshot[29]; CC BY-NC 4.0 |
| Janelia hemibrain | 3D TIN meshes; 3D Tiles, Parquet; Neuroglancer precomputed (990-neuron subset) | *Drosophila* central brain | 25,000 neuron surfaces (the largest traced bodies) | Janelia FlyEM, hemibrain v1.2.1[1]; CC BY 4.0 |
| HuBMAP HRA male body | 3D meshes | Human, whole body | 761 anatomical structures across 24 organs | HuBMAP Human Reference Atlas, CCF v1.3[27]; CC BY 4.0 |
| HuBMAP HRA female body | 3D meshes | Human, whole body | 795 anatomical structures across 31 organs | HuBMAP Human Reference Atlas, CCF v1.3[27]; CC BY 4.0 |

### Extended Data Figs 1 and 2: Polygon level of detail in 2D and Three.js reference viewer on OGC 3D Tiles output

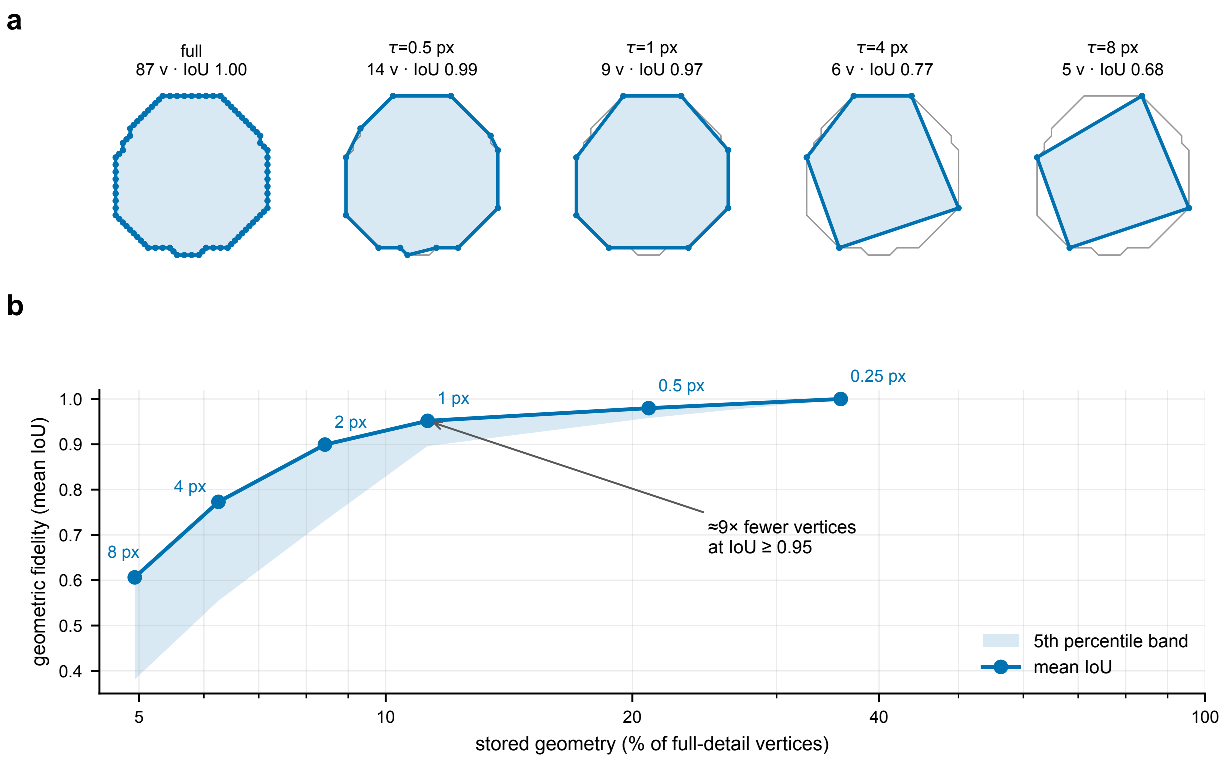


**Extended Data Fig. 1 Polygon level of detail in 2D for PanNuke nuclei. a**, One representative nucleus outline (87 vertices) under progressively larger Douglas–Peucker tolerance $\tau$ (pixels); gray shows the full-resolution boundary and blue indicates the simplified polygon, reduced to a minimum-vertex floor. Vertex count (including the ring's closing vertex) and intersection-over-union (IoU) relative to full detail appear above each example. **b**, Geometric fidelity versus storage across 38,078 nuclei: mean IoU relative to retained-vertex storage (percentage of full-detail vertices, logarithmic scale), with one point per tolerance; shading reaches the 5th-percentile IoU. At 1-pixel tolerance, mean IoU is 0.95 while approximately 11% of vertices are retained (roughly 9× fewer); the initial level eliminates the per-pixel contour staircase at IoU 1.00. Benefits increase with source-vertex detail and are more limited for boundaries already coarse, such as Xenium cells.

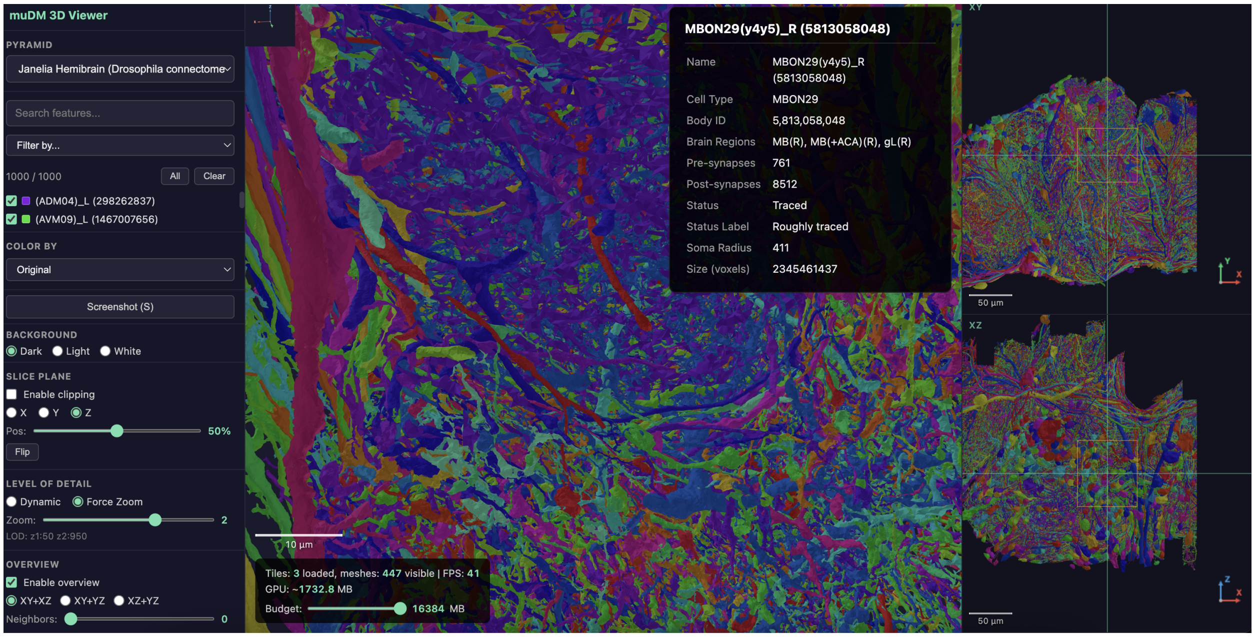


**Extended Data Fig. 2  Three.js reference viewer on OGC 3D Tiles output.** A 1,000-neuron subset of the catalog's 25,000-neuron *Drosophila* hemibrain scene. Left: a feature-selection interface filters neurons by cell type, instance, body identifier, and synapse counts stored in tile metadata. Center: streamed level-of-detail tiles with neuron-specific coloring. Lower left of the center panel: a live display reports loaded tiles, visible meshes, and the bounded GPU-memory budget controlling tile eviction. Far right: XY and XZ overview projections.
